# Visual function restoration in genetically blind mice via endogenous cellular reprogramming

**DOI:** 10.1101/2020.04.08.030981

**Authors:** Xin Fu, Jie Zhu, Yaou Duan, Gen Li, Huimin Cai, Lianghong Zheng, Hao Qian, Changjun Zhang, Zibing Jin, Xiang-Dong Fu, Kang Zhang

**Affiliations:** Center for Biomedicine and Innovations, Faculty of Medicine, Macau University of Science and Technology, Macao, China; Institute for Genomic Medicine and Shiley Eye Institute, University of California San Diego, La Jolla, CA 92093, USA; Guangzhou Women and Children’s Medical Center, Guangzhou Medical University, Guangzhou 510623, China; Department of Nano Engineering, University of California San Diego, La Jolla, CA 92093, USA; Guangzhou KangRui Biological Pharmaceutical Technology Company, 510005 Guangzhou, China; Molecular and Cellular Medicine and Institute for Genomic Medicine, University of California, San Diego, La Jolla, California 92093, USA; Laboratory for Stem Cell & Retinal Regeneration, Institute of Stem Cell Research; Division of Ophthalmic Genetics, The Eye Hospital, Wenzhou Medical University, Wenzhou, 325027 China

**Author notes:** Editorial correspondence: Dr. Kang Zhang, Macau University of Science and Technology, Macao, China. These authors contributed equally to this work (Xin Fu, Jie Zhu).

## Abstract

In this study, we developed an in-situ cellular reprogramming strategy for potent restoration of vision in advanced/end-stage retinitis pigmentosa (RP). Via repressing PTB, an RNA binding protein critical for converting non-neuronal cells to the neuronal lineage, we successfully reprogramed Müller glia to a retinal neuronal fate, and then to cones. We demonstrated that this cellular reprogramming approach was able to rescue retinal photoreceptor degeneration and restore visual functions in two RP mouse models with total blindness, suggesting a novel universal strategy for treating end-stage degenerative diseases.

## Main

The degeneration of retinal neurons is the end point of the most common causes of irreversible blindness, affecting over 50 million people world-wide. Retinitis pigmentosa (RP), a common degenerative eye disease that affects ∼1.5 million people worldwide, is characterized by the progressive degeneration of rod photoreceptors in the retina followed by deterioration and death of cone photoreceptors. As RP patients bear mutations in over 200 causative genes, it is difficult to envision any treatment with conventional gene therapy strategies to correct individual mutations. Therefore, researchers are seeking other cellular sources residing in retina to restore degenerated photoreceptors resulting from the near complete loss of both rods and cones in advanced/end stage RP patients. In non-mammalian vertebrates, such as the zebrafish, retinal injury could be reversed through an intrinsic de-differentiation/reprogramming process by which endogenous Müller glia cells are converted into all retinal neuronal cell types, including photoreceptors, interneurons, and retinal ganglion cells, thereby restoring visual function ^1-4^. However, this regenerative-reprogramming potential is almost non-existent in mammals. Previously, Jorstad *et al*. demonstrated that Müller glia can be reprogrammed into interneurons by combining of engineered expression of Ascl1 and epigenetic modifiers ^5^. A more recent study by Yao *et al.* showed that Müller glia can be reactivated and reprogrammed into rod photoreceptors via sequential gene transfers of ß-catenin and transcription factors essential for rod cell fate specification and determination ^6^. While this approach again demonstrated the reprogramming potential of Müller glia, full vision restoration has not yet been achieved, as only rod photoreceptors were generated. Therefore, despite recent advancements, it has still remained a major challenge in reprogramming müller glia to cone photoreceptors, as the ultimate goal in vision therapy is to restore central precision vision served by cone photoreceptors.

In this study, we were encouraged by our colleague’s results showing a remarkable efficiency in converting astrocytes to functional dopaminergic neurons directly in mouse brain via knockdown of a key RNA binding protein, PTB, which has been previously established as a key gatekeeper for neuronal induction in mammals ^7,8^ (Xiang-Dong Fu *et all*, April 7, 2020 at BioRXiv). This work exploited a unique characteristics of astrocytes in which the neuronal induction is prevented by high-level PTB, but many critical factors for neuronal maturation are expressed, and as a result, transient PTB knockdown alone was sufficient to induce a permanent switch of astrocytes to functional neurons. As Müller cells are the major population of glia in the retinal system, we explored Müller glia for retinal photoreceptor regeneration by using this new approach. Besides using specific photoreceptor cell markers to demonstrate successful conversion, we focused on the determination of cone function and visual acuity by measuring electroretinography (ERG) responses and tested optic kinetic nystagmus (OKN).

We engineered two adeno-associated viral (AAV) vectors for specific PTB inhibition in Müller glia and lineage tracing: AAV-LoxP-Stop-LoxP-RFP-shPTB or AAV-LoxP-Stop-LoxP-RFP, together with AAV-GFAP-Cre-GFP (Fig. 1). Given that the GFAP promoter is only active in astrocytes, including Müller glia, targeted cells with PTB repression could be traced by RFP expression (Fig. 1A). To assess the therapeutic efficiency of this approach in advanced/late stage RP, mice carrying Rd10 mutations were intervened with treatment at postnatal day 90 (P90), in which all rod and cone photoreceptors have degenerated, resulting in complete blindness. Tests for visual function were performed at postnatal day 130, followed by tissue immunofluorescent analysis of cone markers, including cone arrestin (mCAR) and medium wavelength opsin (M-opsin), as well as retinal neuronal markers, including Cone-rod homeobox protein (Crx) and Paired box protein 6 (Pax6) (Fig. 1B).

**Figure 1.**
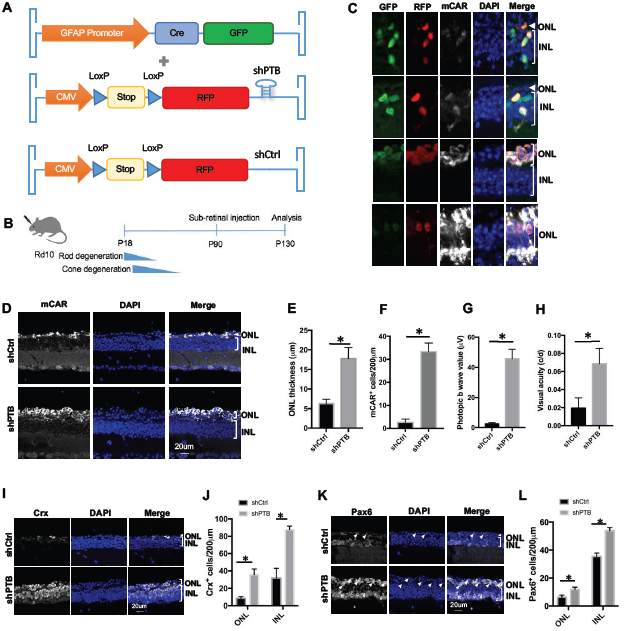
Knockdown of PTB reboots retinal function in 3-month Rd10 mice. (A) Schematics of PTB repression by shRNA in the retina. (B) Experimental scheme for virus injection in Rd10 mice. Mice were treated at P90 and analyzed at P130. Rod degeneration starts around P18, followed by cone degeneration a few days later. No rod and minimal cone activity is detected by P60. (C) Representative image of reprogrammed mCAR^+^ cone photoreceptor from Müller glia in Rd10 mice treated with AAV-shPTB. GFAP-GFP, green; RFP, red; mCAR, grey; DAPI, blue. (D) Immunofluorescent analysis of mCAR^+^ cells in Rd10 mouse retina treated with AAV-shPTB or AAV-shCtrl. mCAR, grey; DAPI, blue. (E) Quantification of mCAR^+^ cells in Rd10 mouse retina treated with AAV-shPTB or AAV-shCtrl. (F) Increased ONL thickness in AAV-shPTB or AAV-shCtrl injected Rd10 mice. ONL, outer nuclear layer. (G) Quantification of b-wave amplitude in AAV-shPTB or AAV-shCtrl injected Rd10 mice (n=6). (H) Quantification of visual acuity in Rd10 eyes treated with AAV-shPTB or AAV-shCtrl (n=6). (I) Immunofluorescent analysis of Crx^+^ cells in Rd10 mouse retina treated with AAV-shPTB or AAV-shCtrl. Crx, grey; DAPI, blue. (J) Quantification of Crx cells in Rd10 mouse retina treated with AAV-shPTB or AAV-shCtrl. Results are shown as mean ± s.e.m. (*p<0.05, student t-test). (K) Immunostaining of retina photoreceptor marker Pax6^+^ in Rd10 mice treated with AAV-shPTB or AAV-shCtrl. Arrows indicated Pax6 expression cells in ONL. Pax6, grey; DAPI, blue. (L) Quantification of Pax6^+^ cells in Rd10 mouse retina treated with AAV-shPTB or AAV-shCtrl. All results are shown as mean ± s.e.m. (*p<0.05, student t-test).

Using this experimental strategy, we found that following PTB repression, Müller glia gradually migrated from inner nuclear layer (INL) to outer nuclear layer (ONL). During this process, cells lost expression of GFP and gained expression of mCAR, indicating a programming process of müller glia to cone photoreceptors (Fig. 1C). To further demonstrate reprogramming of Müller glia in the eye, we performed lineage tracing and following the reprogramming effect of GFAP-marked Müller glia to mCAR-positive photoreceptors in GFAP-Cre wildtype mice. Both rod and cone cells were observed in AAV-shPTB treated retinas (Fig. S1).

We then accessed the visual function in AAV-shPTB treated Rd10 mice. We found that ONL became much thicker and the number of mCAR and Opsin expressing cone-like cells was markedly increased, compared to control mice (Fig. 1D-F, S2-3). We also observed concomitant improvement of visual function as evidenced by a large increase in photopic b-wave value and visual acuity (Fig. 1G-H, S4). These results demonstrated successful induction of functional cone-like cells in mice treated with AAV-shPTB *in vivo.* Meanwhile, increased numbers of positively labeled cells for retinal neuronal markers Crx and Pax6 in both ONL and INL were observed (Fig. 1I-L), consistent with the notion that Müller-to-cone conversion went through an intermediate state for neuronal fate switch. These findings suggest that PTB knockdown leads to the induction of specific transcription factors that are explicitly expressed in photoreceptors, similar to the observation made in the brain.

As mentioned earlier, more than 200 genes are subjected to mutations causing RP. Therefore, to demonstrate the general applicability of our strategy in a distinct mutagenic background, we tested the rescue effect of PTB knockdown in another rapid photoreceptor degeneration mouse model, FVB/N, carrying Rd1 mutations in *pde6b* gene ^9^. Consistent with results from Rd10 mice, treatment of FVB/N retinas with AAV-shPTB showed greater preservation of mCAR^+^ cells and significantly improved preservation of ONL thickness (Fig. 2A-D), along with marked improvement of photopic b-wave values and visual acuity (Fig. 2E-F). Together, these findings demonstrate the reprogramming potential of Müller glia to cone-like cells to account for functional vision restoration in genetically blind mice.

**Figure 2.**
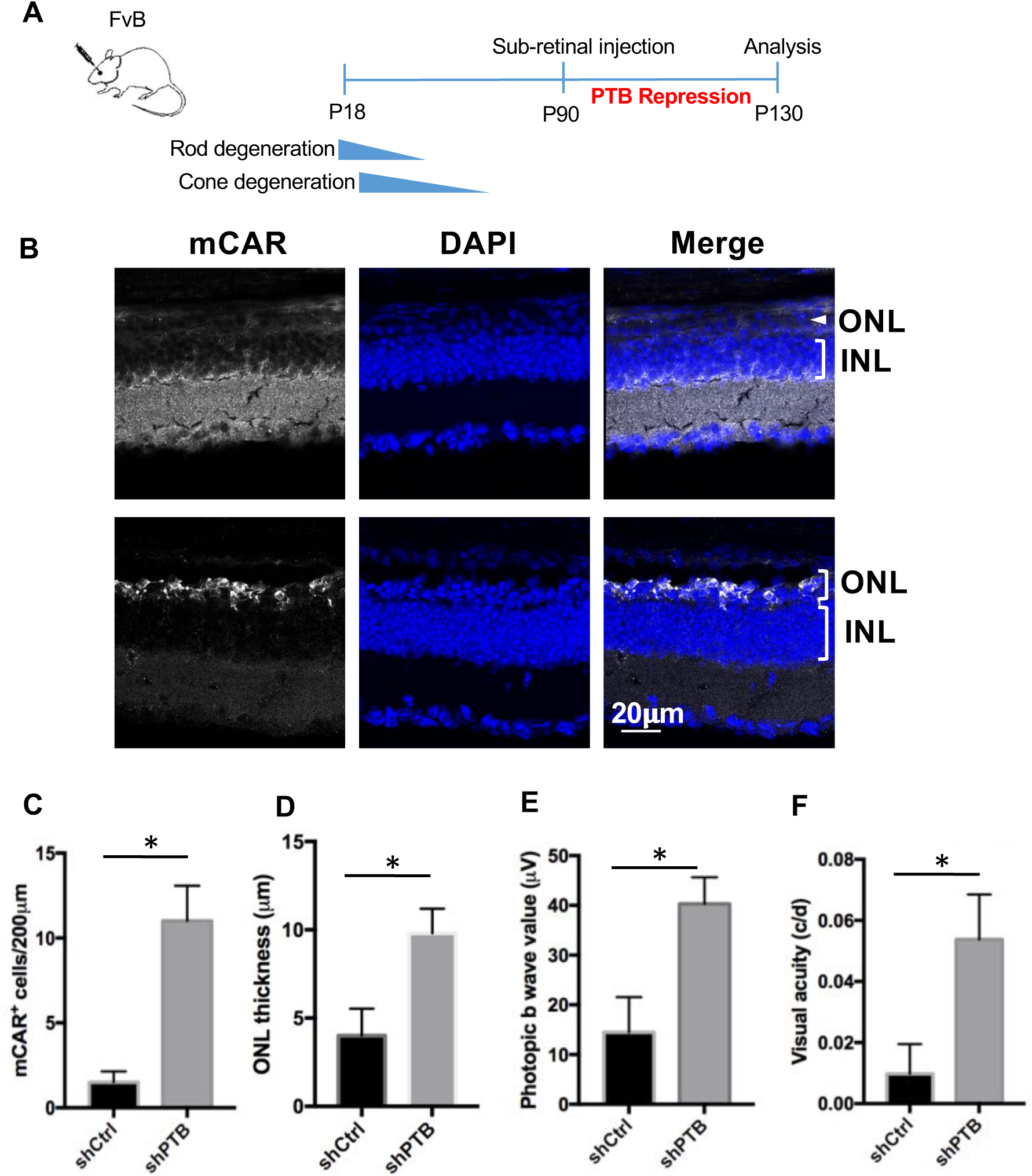
Knockdown of PTB reboots retinal function in 3-month FvB mice. (A) Experimental scheme for virus injection in adult FvB mice. Mice were treated at P90 and analyzed at P130. (B) Immunofluorescent analysis of mCAR^+^ cells in FvB mouse retina treated with AAV-shPTB or AAV-shCtrl. mCAR, grey; DAPI, blue. (C) Quantification of mCAR^+^ cells in FvB mouse retina treated with AAV-shPTB or AAV-shCtrl. (D) Increased ONL thickness in AAV-shPTB or AAV-shCtrl. ONL, outer nuclear layer. (E) Quantification of b wave amplitude in AAV-shPTB or AAV-shCtrl injected FvB mice (n=3). (F) Quantification of visual acuity in FvB eyes injected with AAV-shPTB or AAV-shCtrl. All results are shown as mean ± s.e.m. (*p<0.05, student t-test).

In summary, we exploited a new trans-differentiation strategy by switching Müller glia to cone photoreceptor by depleting a general neuronal induction gatekeeper PTB leading to restoration of visual function in mice with total blindness. These findings suggest a novel universal strategy for treating end-stage retinal degenerative diseases.

## Acknowledgements

This study was funded by Richard Annesser Fund, Michael Martin Fund, Dick and Carol Hertzberg Fund and Maco University of Science and Techonoly.

## Author contributions

X.F., J.Z., Y.D., G.L., L.Z., C.Z., and Z.J. performed mouse *in-vivo* experiments and functional analysis, X.F. and Y.D. performed gene editing work, X.F., Y.D., and Q.H. performed vector construction and AAV production, X.F., J.Z., H.C., and X.S analyzed data. K.Z designed the study. K.Z. X.F. and XD.F wrote the paper. All authors discussed the results and commented on the manuscript.

## Methods

### AAV plasmids and viral production

Plasmids pAAV-LoxP-Stop-LoxP-RFP-shPTB (shPTB) and pAAV-LoxP-Stop-LoxP-RFP (shCtrl) were kind gifts from Xiang-Dong Fu’s lab. AAV-shPTB and AAV-shCtrl with serotype 8 were generated in house utilizing a protocol published previously^1^. Briefly, AAV2/8 virus particles were produced using HEK293T cells via the triple transfection method and purified via SP column chromatography. Confluency at transfection was about 70%, and each plate was transfected with 4ug of pXR8, 4ug recombinant transfer vector, and 5.5ug of pHelper vector using polyethylenimine (PEI; 1 mg/mL linear PEI in DPBS [pH 4.5], using HCl) at a PEI: DNA mass ratio of 4:1. The mixture was incubated for 10 min at room temperature and then applied dropwise onto the media. The virus was harvested after 72 hours and purified via SP column chromatography. The virus was dialyzed with PBS supplemented with 0.05% of Pluronic F68 (Thermo Fisher Scientific) using 100-kDa filters (Millipore). The titer of virus was quantified by qPCR by primers specifically to CMV promoter. AAV-GFAP-GFP-Cre was purchased from University of Pennsylvania Vector Core.

### Animals

All strains including wild type (C57BL/6), Rd10, FvB, FvB-GFAP-GFP (FvB/N-Tg(GFAP-GFP)14Mes/J), and GFAP-Cre (B6.Cg-Tg(Gfap-cre)77.6Mvs/2J) were purchased from the Jackson Laboratory. All mice used in this study were from mixed gender, P1 to 12 weeks old. All mouse experiments were approved by the IACUC committee. All procedures were conducted with the approval and under the supervision of the Institutional Animal Care Committee at the University of California San Diego and adhered to the ARVO Statement for the Use of Animals in Ophthalmic and Vision Research.

### Subretinal injection in mice

Rd10, FvB, FvB-GFAP-GFP, and GFAP-Cre mouse were used in the study. Subretinal injection into P7 neonate and 3-month adult eyes was performed as previously described ^2,3^. Approximately 0.5µl AAV8 (∼2×10^10^GC) were introduced into the subretinal space using a pulled angled glass pipette controlled by a FemtoJet (Eppendorf). Adult mice were anesthetized with an intraperitoneal injection of a mixture of ketamine and xylazine, and neonatal mice were anesthetized by induction of hypothermia. Pupils were dilated with 1% topical tropicamide. The subretinal injection was performed under direct visualization using a dissecting microscope with a pump microinjection apparatus (Picospritzer III; Parker Hannifin Corporation) and a glass micropipette (internal diameter 50∼75 µm). 0.5 µl of AAV mixture was injected into the subretinal space through a small scleral incision. A successful injection was judged by the creation of a small subretinal fluid bleb. Mice showing any sign of retinal damage such as bleeding were discarded and excluded from the final animal counts.

### ERG recording

To monitor the efficacy of gene knock-out in Rod/Cone fate switch, ERG studies were performed at 40 days, 3 months or 6 months after treatment before the animals were sacrificed for histology. Rd10 mice were deeply anesthetized as described for the surgical procedure above. Eyes were treated with 1% topical tropicamide to facilitate pupillary dilation. Each mouse was tested in a fixed state and maneuvered into position for examination within a Ganzfeld bowl (Diagnosys LLC). One active lens electrode was placed on each cornea, with a subcutaneously placed ground needle electrode positioned in the tail and the reference electrodes placed subcutaneously in the head region approximately between the two eyes. Light stimulations were delivered with a xenon lamp in a Ganzfeld bowl. The recordings were processed using software supplied by Diagnosys. Photopic ERG was performed according to a published protocol. Mice were light adapted for 10 minutes at a background light of 30 cd/m^2^. Cone responses were elicited by a 34 cds/m^2^ flash light with a low background light of 10 cd/m^2^, and signals were averaged from 50 sweeps.

### Optokinetic test

Visual acuity testing of all animals was conducted at 5 weeks after injection with an optomotor testing apparatus as previously reported ^4,5^. Briefly, a virtual reality chamber was created with four computer monitors facing into a square. A virtual cylinder, covered with a vertical sine wave grating, was drawn and projected onto the monitors using software running on a Java application. The animal was placed on a platform within a transparent cylinder (diameter ∼30 cm) in the center of the square. A video camera situated above the animal provided real-time video feedback on another computer screen. From the mouse’s point of view, the monitors appeared as large windows through which the animals viewed the rotating cylinder. Each mouse was placed on the platform in a quiet environment before the test until it became accustomed to the test conditions with minimal movement. The virtual stripe cylinder was set up at the highest level of contrast (100%, black 0, white 255, illuminated from above 250 cd/m^2^), with the number of stripes starting from 4 per screen (2 black and 2 white). The test began with 1 min of clockwise rotation at a speed of 13. (The baseline value is 10, at which the bars move 1 pixel/cycle. Values less than 10 delay the cycle by X*100 ms, with a minimum value of 1) An unbiased observer tracked and recorded the head movements of the mouse. The test was then repeated with 1 min of counterclockwise rotation. The data were measured by cycles/degree (c/d) and expressed as mean ± s.e.m., with comparison using a t-test statistical analysis. A p value <0.05 was considered statistically significant.

### Histological analysis of the mouse eye

Following ERG recordings, mice were sacrificed, and retinal cross-sections were prepared for histological evaluation of ONL preservation. mice were euthanized with CO_2_, and eyeballs were dissected out and fixed in 4% PFA. Cornea, lens, and vitreous were removed from each eye without disturbing the retina. The remaining retina containing eyecup was infiltrated with 30% sucrose and embedded in OCT compound. Horizontal frozen sections were cut on a cryostat. Retinal cross-sections were prepared for histological evaluation by immunofluorescence staining.

### Immunofluorescence

Retinal cryosections were rinsed in PBS and blocked in 0.5% Triton X-100 and 5% BSA in PBS for 1 hour at room temperature, followed by an overnight incubation in primary antibodies at 4°C. After three washes in PBS, sections were incubated with secondary antibody. Cell nuclei were counterstained with DAPI (49,6-diamidino-2-phenylindole). The following antibodies were used: mouse anti-Rhodopsin monoclonal antibody (Abcam ab3267), rabbit anti-Cone Arrestin polyclonal antibody (Millipore AB15282), rabbit anti-Opsin polyclonal antibody (Millipore AB5405), Mouse anti-Crx (Abnova, H00001406-M02), Rabbit anti-Pax6 (Biolegend, 901301). The secondary antibodies, Alexa Fluor-488- or 555 or 647-conjugated anti-mouse or rabbit or chicken immunoglobulin G (IgG) (Invitrogen) were used at a dilution of 1:500. Sections were mounted with Fluoromount-G (Southern Biotech) and coverslipped. Images were captured using an Olympus FV1000 confocal microscope.

### Statistical analysis

Results are shown as mean ± s.e.m., as indicated in the figure legend. Comparisons were performed with student’s *t*-test, as indicated in the figure legend.

## Supplementary figure legends

**Figure S1.**
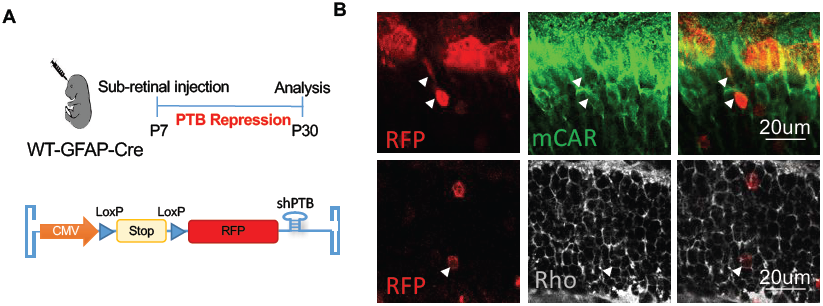
Reprogramming of Müller glia into rods and cones via PTB repression. (A) Experimental scheme for virus injection in newborn GFAP-Cre mice. Mice were treated at P7 and analyzed at P30. (B) Immunofluorescent analysis of mCAR and Rhodopsin (Rho) in GFAP-Cre mice. GFAP-Cre (B6.Cg-Tg(Gfap-cre)77.6Mvs/2J) were subretinally injected with AAV-shPTB. Müller glia was traced by RFP when PTB was knocked down. Arrows indicate reprogrammed rod or cone cells from GFAP expression müller glia. mCAR, green; RFP, Red; Rhodopsin, grey.

**Figure S2.**
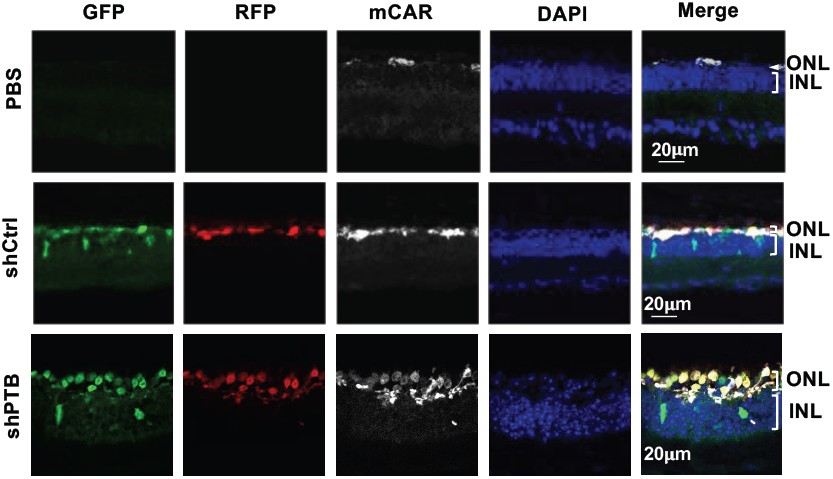
Knockdown of PTB significantly increased mCAR^+^ cells in 3-month Rd10 mice. Representative images of immunofluorescent analysis of mCAR^+^ cells in Rd10 mouse retina treated with AAV-shPTB, AAV-shCtrl, or PBS. Mice were treated at P90 and analyzed at P130. GFAP-GFP, green; RFP, red; mCAR, grey; DAPI, blue.

**Figure S3.**
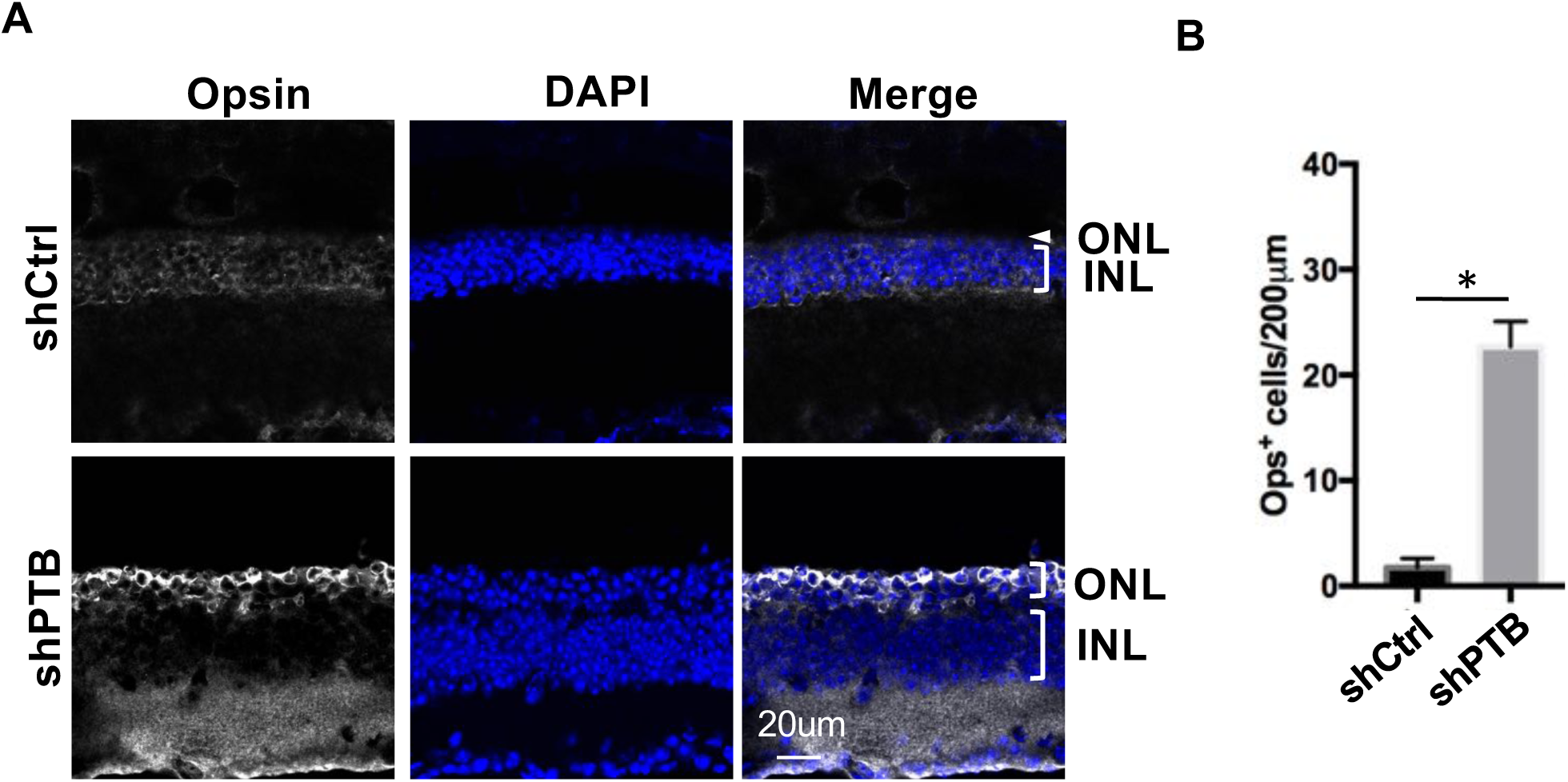
Knockdown of PTB significantly increased Opsin^+^ cells in 3-month Rd10 mice. (A) Immunofluorescent analysis of Opsin^+^ cells in Rd10 mouse treated with AAV-shPTB or AAV-shCtrl. Rd10 mice were treated at P90 and analyzed at P130. Opsin, grey; DAPI, blue. (B) Quantification of Opsin^+^ cells in Rd10 mouse retina treated with AAV-shPTB or AAV-shCtrl. Results are shown as mean ± s.e.m. (*p<0.05, student t-test).

**Figure S4.**
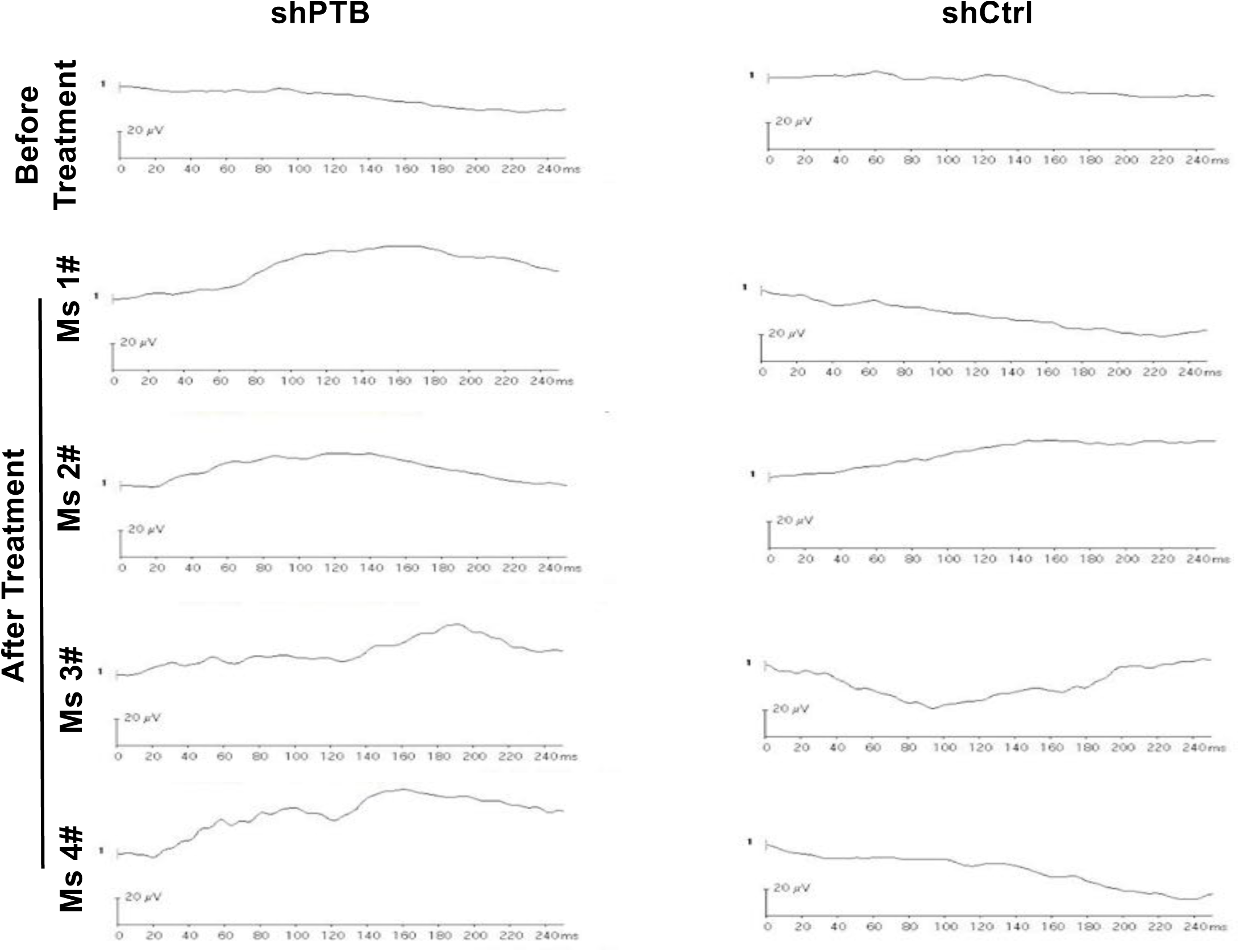
Representative images of Photopic ERG recording in Rd10 retina treated with AAV-shPTB or AAV-shCtrl.

